# Drivers of diversity in individual life courses: Sensitivity of the population entropy of a Markov chain

**DOI:** 10.1101/188276

**Authors:** Ulrich K. Steiner, Shripad Tuljapurkar

**Affiliations:** Center for Research and Interdisciplinarity, Paris, France; Department of Biology, University of Southern Denmark, Odense, Denmark; Department of Biology, Stanford University, USA

## Abstract

Individuals differ in their life courses, but how this diversity is generated, how it has evolved and how maintained is less understood. However, this understanding is crucial to comprehend evolutionary and ecological population dynamics. In structured populations, individual life courses represent sequences of stages that end in death. These sequences can be described by a Markov chain and individuals diversify over the course of their lives by transitioning through diverse discrete stages. The rate at which stage sequences diversify with age can be quantified by the population entropy of a Markov chain. Here, we derive sensitivities of the population entropy of a Markov chain to identify which stage transitions generate—or contribute—most to diversification in stage sequences, i.e. life courses. We then use these sensitivities to reveal potential selective forces on the dynamics of life courses. To do so we correlated the sensitivity of each matrix element (stage transition) with respect to the population entropy, to its sensitivity with respect to fitness λ, the population growth rate. Positive correlation between the two sensitivities would suggest that the stage transitions that selection has acted most strongly on (sensitivities with respect to λ) are also those that contributed most to the diversification of life courses. Using an illustrative example on a seabird population, the Thick-billed Murres on Coats Island, that is structured by reproductive stages, we show that the most influential stage transitions for diversification of life courses are not correlated with the most influential transitions for population growth. Our finding suggests that observed diversification in life courses is neutral rather than adaptive. We are at an early stage of understanding how individual level dynamics shape ecological and evolutionary dynamics, and many discoveries await.

## Introduction

In any population we observe great diversity in phenotypes and life courses among individuals (Tuljapurkar et al., 2009; Steiner and Tuljapurkar, 2012). How such diversity is generated, how it has evolved and how maintained is of interest to population biologists, biodemographers, evolutionary biologists, and ecologists, because such knowledge furthers understanding of ecological and evolutionary change (Endler, 1986; Hartl and Clark, 2007). This interest has propelled analyses of how genetic variability, environmental variability and their interaction generate individual differences in phenotypes and life courses. Population genetic models focus on mutations, drift, and so on to explain genotype frequencies and their dynamics (Hartl and Clark, 2007; Barton and Keightley, 2002; Mackay et al., 2009; Orr, 2005; Der et al., 2011). A challenge not fully mastered, is how these mechanisms lead to stable populations that show the kind of variability observed in natural populations (Evans and Steinsaltz, 2007; Roze and Rousset, 2008). Quantitative genetics circumvents some of these challenges by investigating phenotypic trait distributions and their changes within populations (Walsh, 2001; Barton et al., 2017). Environmental variation leads to changes in the phenotype, and genotype-environment interaction further adds to the complexity in understanding observed diversity in phenotypes and life courses (Champagnat et al., 2006). Phenotypic plasticity investigates these genotype-environment interactions, and processes such as niche construction and eco-evolutionary feedback emphasize that the population’s environment is not fixed, but interacts with and can be altered by the organism (Diekmann et al., 2003; Vuilleumier et al., 2010; Pelletier et al., 2009). Ideas about neutral variability and epigenetics have also been used to explain the observed diversity of genotypes, phenotypes, and life histories (Ohta and Gillespie, 1996; Steiner and Tuljapurkar, 2012; Geoghegan and Spencer, 2012). Neutral concepts include non-adaptive phenotypic variation due, e.g., to spandrels—phenotypes as byproducts of selection on other traits, or genetic hitchhiking (Evans and Steinsaltz, 2007; GOULD and LEWONTIN, 1979). Most of the above concepts are considered to be generally applicable across biological systems. However, these concepts are challenged to explain the surprising diversity in life courses of even isoclonal individuals raised under highly controlled environmental conditions (Lande et al., 2003; Finch and Kirkwood, 2000; Melbourne and Hastings, 2008; Steiner and Tuljapurkar, 2012; Jouvet et al., 2018; Steiner et al., 2019). The challenges arise because these concepts do not consider the underlying individual level dynamics that contribute substantially to the diversity in individual life courses. Besides the lack of understanding of drivers of individual level dynamics, we often do not know to what degree these drivers are adaptive, maladaptive or neutral (Lenormand et al., 2009).

Whatever the actual mechanisms may be, the diversity in life courses in any structured population can be characterized by differences among stage trajectories—sequences of stages that individuals go through over their life course and that end in death (Caswell, 2001; Tuljapurkar et al., 2009). Here we assume stages are discrete (this is just binning). Individuals are born into one, or one of several, discrete stages and subsequently transition to one of several discrete stages at each observation. If the transition probability only depends on the current stage, these trajectories can be described by a Markov chain. Over *L* observations—think of one observation per year—with *s* stages there are a maximum of *s*^*L*^ possible trajectories, i.e. trajectories diversify with increasing length *L*. The larger the uncertainty at each step, the larger is the diversity of life course trajectories (Tuljapurkar et al., 2009). Stages include developmental stages including levels of breeding success, morphological stages such as size, behavioral stages such as feeding or mating activity, physiological stages such as condition, gene expression stages such as transcription factor expression, epigenetic stages such as methylation stage, or spatial location.

In this paper, individual trajectories are described by a Markov chain, i.e., there is a probability *p*_*ij*_ ≥ 0 that an individual changes its stage from stage *j* to stage *i*, for every possible pair of stages. The notation here is similar to Caswell (2001); Hill et al. (2004). In many systems the stage distribution at birth is centered on one or a few stages. With increasing age, individuals transition through stages described by the Markov chain and individual stage trajectories diversify. We can quantify the rate of diversification of these trajectories by the entropy of the Markov chain (Shannon, 1948). This entropy has been termed population entropy (Tuljapurkar et al., 2009). The process of diversification of life courses by Markovian (stochastic) stage transitions has been called dynamic heterogeneity with its outcome of individual differences (Tuljapurkar et al., 2009; Steiner and Tuljapurkar, 2012; Caswell, 2009). This process, based on transitions with identical probabilities but different outcomes, contrasts with fixed differences in transition rates. With fixed differences, each genotype is described by its own matrix of transition rates.

Here we focus on the sensitivity of population entropy to the underlying set of transition probabilities. These sensitivities should reveal which transitions generate the most diversification among life courses. These sensitivities of the population entropy, however do not provide any understanding whether such diversification might be under selection, i.e. whether it is adaptive, maladaptive, or neutral. To investigate potential adaptive features, we conider each transition rate and compare the sensitivity of the population entropy to the sensitivity of the population growth rate, λ. This latter sensitivity to λ is linked to the evolutionary forces acting on these transition probabilities, because population growth rate quantifies fitness (Caswell, 2001). A positive correlation between sensitivities suggests that diversification should be adaptive; diversification is neutral if we do not see any relationship between the sensitivities; and diversification may be maladaptive if the sensitivities are negatively correlated.

We describe sensitivities for ergodic Markov chains, and Markov chains with absorbing stages. In most applied cases the absorbing stage is the death stage. Classical population projection matrix models that include reproduction (e.g. Lefkovitch or Leslie population matrix models) first need to be transformed into a Markov chain before we can estimate the population entropy. We can achieve this transformation as described by Tuljapurkar (1982) (Appendix). We illustrate our results for a seabird population, the Thick-billed Murre on Coats Island, Canada (Gaston et al., 1994; Steiner and Gaston, 2005). This population is structured by reproductive stages, defined as breeding outcomes.

Our results have the virtue that they only require the dominant eigenvalue and corresponding eigenvectors of non-negative matrices—these are numerically straightforward and well-conditioned, unlike the computation of all subdominant eigenvalues. Our approach is therefore applicable to many structured populations.

### Population entropy and Matrix of a Markov chain

When the population is ergodic (actually, irreducible and aperiodic) there is a stationary (or equivalently, equilibrium) frequency distribution over the possible stages: a vector **w** whose elements *w*_*i*_ are the frequencies of stages *i* = 1, *…*, *s*. A stage’s equilibrium frequency also equals the fraction of times that an individual is expected to be in that stage, if we make many repeated observations. Population entropy *H*(**P**) quantifies the diversity in individual trajectories described by the Markov chain:

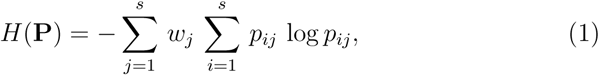

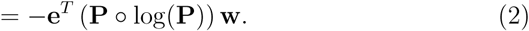

Here **P** is a matrix of the Markov chain transition probabilities *p*_*ij*_, with individuals transitioning from column *j* to row *i*. The second line above is useful numerically and analytically: the superscript *T* indicates a transpose; **e** is a vector whose entries all equal 1; the Hadamard product (∘) is elementwise so that for matrices **P**, log(**P**) of equal size with elements *p*_*ij*_, *log*(*p*_*ij*_) respectively the matrix **P** ∘ log(**P**) is of same size and has *ij* element equal to *p*_*ij*_ *log*(*p*_*ij*_).

We start with deriving sensitivities for an ergodic chain (irreducible, non-absorbing), by asking what happens if we make a small change in the transition probabilities so that **P** becomes **P** + *ϵ* **B** (for small positive *ϵ*). Throughout this paper, we consider only perturbations that leave unchanged the signature of the Markov chain: i.e., whenever *p*_*ij*_ = 0 we keep *b*_*ij*_ = 0. Then the population entropy must change from *H*(**P**) to say *H*(**P**) + *ϵ H*_1_. Then *H*_1_ is the sensitivity of the population entropy. We obtain here an exact analytical expression for this sensitivity.

Thereafter, we answer the analogous question for a Markov chain that has at least one “absorbing” stage. To see why this is different, suppose death is the absorbing stage so that an individual wanders among the non-absorbing stages until it dies. Conditional on being alive, we expect that there is a quasi-stationary distribution over the non-absorbing stages, if we can find appropriate conditional Markov transition probabilities. Darroch and Seneta (1967) show that we can, providing that absorption takes a long time; see also Matthews (1970). The entropy of this conditional Markov chain measures the rate of individual trajectory diversification until death. Our contributions are an exact result for the sensitivity of the population entropy of an ergodic chain and absorbing Markov chains. Comparing the sensitivities between the two types of Markov chains (ergodic and absorbing) from the same system can then be used to evaluate the contribution of individuals surviving to different ages on the diversity of stage trajectories, as has been done before (Hernandez-Pacheco and Steiner, 2017).

## Sensitivity of Entropy: Ergodic Chains

### Changing Transition Probabilities

The starting point is a population described by a matrix **P** of transition probabilities; we assume the chain is irreducible and aperiodic, hence ergodic. An ergodic population is characterized by its asymptotic dynamics being independent of the starting conditions. Here, we are mainly interested in such ergodicity since our focus is on revealing underlying processes, i.e. the drivers of diversity in life courses, than on initial conditions a population starts at. For such ergodic models the stationary frequency is an right eigenvector, **Pw** = **w**. Transition probabilities out of each stage sum to unity, so **e**^*T*^ **P** = **e**^*T*^. We compute the fundamental matrix, which has also been described as the stage duration matrix (Steiner et al., 2012).

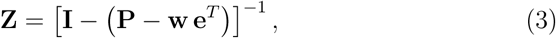

where **I** is the identity matrix, and ^−1^ indicates the inverse of the function.

Now perturb the transition probabilities to **P**+ *ϵ* **B**, so that transition probability *p*_*ij*_ changes to *p*_*ij*_ + *ϵb*_*ij*_. Clearly we must have

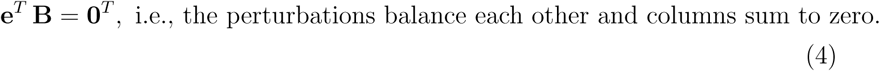

This means that changes in the transition probabilities are necessarily constrained, we cannot simply perturb only a single *p*_*ij*_; some biologically distinct ways of achieving this constraint are discussed by Caswell (2001), pages 218-220.

Following Schweitzer (1968) the stationary frequencies change to **w**+ *ϵ* **y** + *ϵ*^2^**y**_2_ + *O*(*ϵ*^3^) where **e**^*T*^ **y** = **e**^*T*^ **y**_2_ = 0

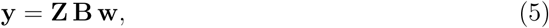

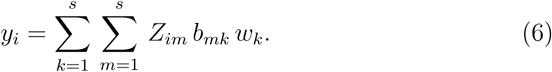

The more involved expression for **y**_2_ is found in Kato (1966). So the vector **y** from equation (5) comprises, first, the time an individual spends in each stage given its current stage (i.e. the fundamental or stage duration matrix, **Z**), second, the product with the perturbation matrix **B** then determines the change in time each individual spends in each stage given its current stage, and finally, the multiplication with the stable stage distribution **w** quantifies how many individuals (or more precisely what proportion of individuals) are affected by the change in time they spent in each stage. That is the final multiplication with the stable stage distribution **w** quantifies how many individuals are affected by how much time they spend in each stage due to the perturbation, which is exactly how much change in the stationary stage distribution is caused by the perturbation.

### Sensitivity of Entropy

From equation (1) (and the Appendix) the entropy of the perturbed Markov chain is

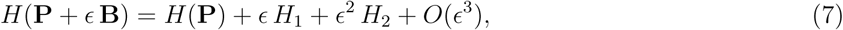

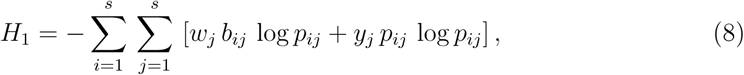

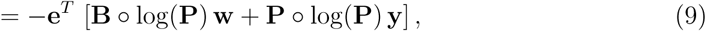

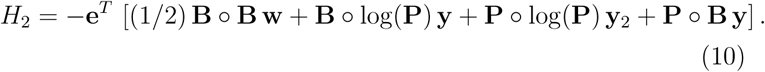

Here *H*_1_ is the sensitivity to the population entropy we seek. The second-order change in entropy (essentially the second derivative) is *H*_2_. For equation (8) we have the stage distribution element *w*_*j*_ (how many individuals are affected), by the amount of perturbation *b*_*ij*_, and the change in stage distribution *y*_*j*_. An illustration for the special case of perturbing a Maximum Entropy chain is given in the Appendix.

## Sensitivity of Entropy: Chains with Absorbing stages

### Transition Probabilities with Absorption

We consider just one absorbing stage—multiple absorbing stages are easily dealt with (Matthews, 1970). Let us say the absorbing stage (think “death”) is the last stage of *s* stages, so that stages 1 to (*s* − 1) are the transient (i.e., “alive”) stages. The transition probability matrix must have the form

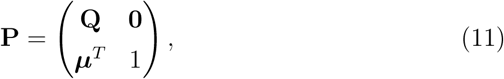

with absorption (death) probabilities given by the elements *µ*_*i*_ of vector ***µ*:**

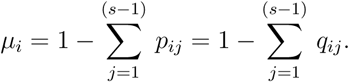

Matrix **Q**, describes the transition probabilities among the life stages, summing over the columns of **Q** gives the survival probability of each stage. Conditional on non-absorption (i.e., being alive), the transition probabilities among the (*s* − 1) transient stages (Darroch and Seneta, 1967) are the entries in the (*s* − 1) × (*s* − 1) matrix

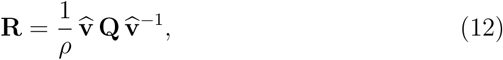

where 0 *< ρ <* 1 is the dominant eigenvalue of **Q, v** with elements *v*_*i*_ is the corresponding left eigenvector,

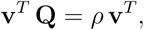

and the diagonal matrix

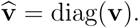

The *ij* element of matrix **R** is *v*_*i*_*q*_*ij*_*/*(*ρv*_*j*_); clearly, the columns of **R** sum to 1, so this is a Markov matrix, while matrix **Q** is not. So what we have done in (12) is to transform the transient (absorbing stage transition) matrix **Q** to a Markov chain **R**. Let **w** be the right eigenvector of **Q** corresponding to its dominant eigenvalue, normalized so that (**v**^*T*^ **w**) = 1. The equilibrium frequency distribution of the conditional process governed by **R** is given by the products (*w*_*i*_*v*_*i*_), *i* = 1 *…* (*s* − 1).

We can measure the diversification of individual trajectories with increasing age while they are still alive by the population entropy of the conditional process (see Appendix),

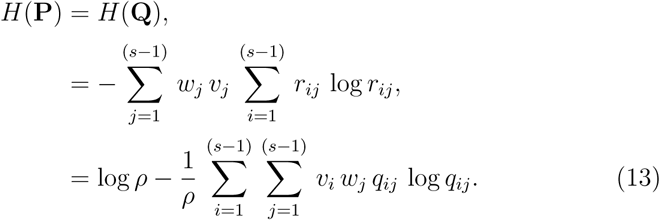

### Perturbing an Absorbing Chain

We now want the effect on the population entropy of small changes in the transition probabilities of the Markov chain. In (11), consider simple changes in the transient matrix **Q** to **Q** + *ϵ* **B**. It is easy to see how this changes the full matrix **P**. These changes will alter *ρ*, **v**, and **w** to *ρ* + *ϵv*, **v** + *ϵ***x, w** + *ϵ***y**, respectively. Here we give explicit formulas to compute these changes and in the next subsection show how these are used to compute the sensitivity of entropy we seek.

Recalling that (**v**^*T*^ **w**) = 1, we have the well-known (see e.g., Caswell (2001)) fact that

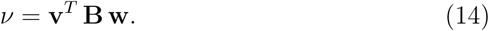

We define two new matrices:

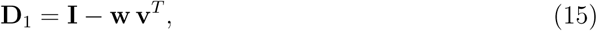

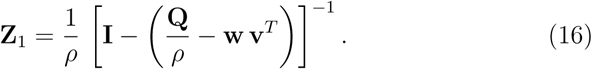

Then we have (see Appendix) the less well-known results,

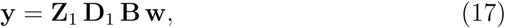

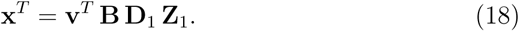

The interpretation of **y** in (17) is similar to the one in equation (5), i.e. how many individuals are affected by how much (more or less) time they spend in each stage due to the perturbation, which equals how much change in the stationary stage distribution is caused by the perturbation, except here (17) this change is based on the absorbing (transient) transition matrix.

### Sensitivity of Entropy for an Absorbing Chain

The last step is to compute the difference between the entropy of the perturbed chain (*H*(**Q**)) and the original chain,

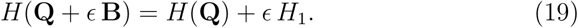

The sensitivity *H*_1_ is given (see Appendix) by

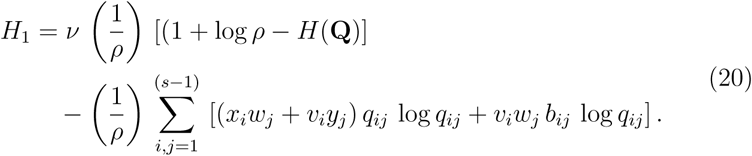

## Illustrative example sensitivity of population entropy: The Thick-billed Murre

To illustrate our exact result for the sensitivity of the population entropy of a Markov chain, we first built a stage-structured matrix population model using longitudinal mark-recapture data on a highly philopatric and colonial seabird species, the Thick-billed Murre *(Uria lomvia)* (Gaston et al., 1994; Steiner and Gaston, 2005). After parameterizing the population projection matrix based on the longitudinal data, we transformed this matrix to a Markov chain, as described by Tuljapurkar (1982) (Appendix, see also equation (12)). Here we present the results on population entropy (ergodic chain) of the resulting Markov chain and discuss its implications.

### Structured population model of the Thick-billed Murre

To parameterize the stage-structured matrix model, we used data on 1984 individual seabirds, Thick-billed Murres, banded between 1981 and 2010, on Coats Island, Nunavut, Canada (62°30′*N*, 83°00′*W*). Band readings have been made between 1991 and 2011 in the colony over each breeding season. For each bird for which a band was read its breeding status (breeding outcome) for that season was recorded as a) I, immature, birds prior to any breeding attempt; b) E, egg laid, bird laid an egg but the egg did not hatch; c) H, hatch, bird managed to hatch a chick but the chick did not fledge; d) F, fledged, the bird’s chick fledged, i.e. chick disappeared *>*=10 days after hatching; or e) U, unknown, when the breeding outcome of the bird was not known. Birds are born into the immature stage (I) and they remain in that stage until they are three years old (only 3 out of the 1128 individuals banded as chicks, i.e. known aged birds, recruited at age two into the breeding cohort). After the third year, individuals can stay as immatures, or transition to and then among one of the other breeding outcome stages, E, H, and F. Since some birds had unknown breeding stages, we corrected the estimated survival and transition probabilities among the observed breeding stages (E, H, F) for the unknown events by weighting probabilities according to survival and transition rates (Appendix).

Our resulting stage structured matrix projection model included the four stages (I,E,H,F), with stage F being the only stage contributing to re-production. Since sex determination for Thick-billed Murres is challenging, we used data on both sexes for estimating survival, recapture (sighting), and transition probabilities (assuming same survival and transitioning for both sexes). We assumed 50% of chicks to be female, and we included only females for the fertility of the projection model (Table 1). Further detail on estimating resighting, survival and transition probabilities, for which we used program MARK (White and Burnham, 1999), is provided in the Appendix. The corresponding transformed Markov chain (see equation (12)) is shown in Table 2.

**Table 1:** Projection matrix model.

|  | I | E | H | F |
| --- | --- | --- | --- | --- |
| I | 0.494 | 0 | 0 | 0.5 |
| E | 0.22 | 0.465 | 0.231 | 0.161 |
| H | 0.006 | 0.116 | 0.094 | 0.088 |
| F | 0.02 | 0.378 | 0.55 | 0.657 |

**Table 2:**
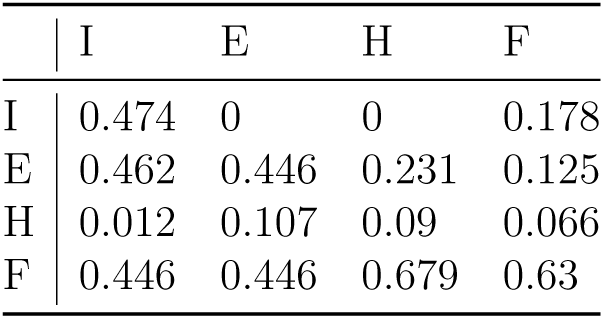
Tranformed Markov chain matrix.

|  | I | E | H | F |
| --- | --- | --- | --- | --- |
| I | 0.474 | 0 | 0 | 0.178 |
| E | 0.462 | 0.446 | 0.231 | 0.125 |
| H | 0.012 | 0.107 | 0.09 | 0.066 |
| F | 0.446 | 0.446 | 0.679 | 0.63 |

### Demographic parameters of the stage structured Thick-billed Murre population model

We estimated the population growth rate for the projection model at λ=1.041 (dominant eigenvalue of matrix shown in Table 1), which might be a slight overestimation compared to the observed population growth; accounting for stochastic environmental variation would lower the expected growth rate slightly. The quasi stable stage distribution of the projection model was I=0.33, E=0.25, H=0.07, F=0.36 (scaled corresponding right eigenvalue **w**) and the corresponding reproductive values are I=1.0, E=2.2, H=2.1, F=2.7 (corresponding left eigenvalue **v**, scaled for I=1). The sensitivities with respect to λ of the population projection model (Table 1) are given in Table 3 and estimated according to Caswell (2001) (page 209ff). They show that population growth rate is most sensitive to transitions from the immature to the fledging stage, as well as remaining in the fledging stage, the only stage that contributes to fertility. Moving from population growth—and its sensitivity—to evaluating diversification, the population shows a high rate of diversification with an population entropy (H=0.98%) close to the maximum entropy for the Markov chain matrix (Table 2).

**Table 3:**
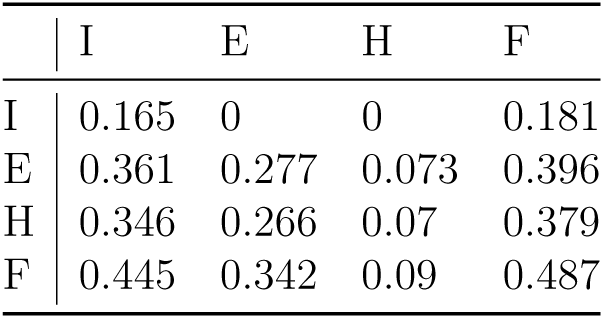
Sensitivity to λ.

|  | I | E | H | F |
| --- | --- | --- | --- | --- |
| I | 0.165 | 0 | 0 | 0.181 |
| E | 0.361 | 0.277 | 0.073 | 0.396 |
| H | 0.346 | 0.266 | 0.07 | 0.379 |
| F | 0.445 | 0.342 | 0.09 | 0.487 |

### Integrated sensitivities and selective forces

The sensitivities with respect to λ of the population projection model, as we estimate for instance in Table 3, imply that a realized perturbation in a transition probability *p*_*ij*_ alters the survival rate of that stage *j*. That is, if we increase a transition rate, *p*_*ij*_, in a given stage *j*, we automatically increase the column sum across transitions in the given stage *j* by the same amount; the column sum determines the survival rate of a stage. Here we are not interested in relationships between reproduction and survival, but in changes among stage dynamics without changing stage survival. We therefore need to keep the column sum of the stage constant when we perturb a transition probability. This constraint implies, if we perturb one transition probability we have to compensate this perturbation by one or more matrix elements in the same column, i.e. transition rates in the same stage. The biological implications of such constraints in changing the transition probabilities for stage structured models are discussed by Caswell (2001) (pages 2018-2019). There are many solutions to fulfill these constraints, here, we reduced (perturbed) the transition probability of one matrix parameter by 0.01 and increased at the same time the transition probabilities of the remaining stage parameters by equal amounts as to perturbations **e**^*T*^ **B** = **0**^*T*^, i.e., columns sum of the perturbations equal zero (see also equation (4)). We call these sensitivities integrated sensitivities following Van Tienderen (1995); these integrated sensitivities comprise changes in multiple transition rates and we sum weighted sensitivities according to the perturbations described in **B**. These constraints on the perturbations (**e**^*T*^ **B** = **0**^*T*^) ascertain the assumption (requirement) of ergodicity of the matrix model (Markov chain). We estimated such an integrated sensitivity related to a reduction in each transition probability (note we consider only perturbations that leave unchanged the signature of the Markov chain: i.e., whenever *p*_*ij*_ = 0 we keep *b*_*ij*_ = 0). Each change in a transition probability changes the population entropy (diversification in life courses) and the population growth (λ), but perturbations now having signs, and resulting changes on population entropy or population growth can be positive or negative. Classical sensitivities, as illustrated for instance in Table 3, hold only positive values; any increase in a transition rate also increases survival and therefore has to increase population growth. Classical sensitivities do not evaluate changes among stage dynamics as we do here.

In Table 4 we show results for the integrated sensitivities of population entropy for the Thick-billed Murre example. Table 5 shows the corresponding integrated sensitivities with respect to λ. If we reduce the transition rate of remaining as immatures (I to I, *b*_1,1_) by 0.01, and at the same time increase the remaining three transition probabilities (from I to E,H & F, *b*_2,1_ to *b*_4,1_) by 0.01*/*3 = 0.003333, population entropy increases by 0.0034 (first element Table 4), while the population growth rate, λ, increases by 0.00219 (first element Table 5). A reduction in the probability of birds successfully fledging a chick in two consecutive years (transition stage F to F, *p*_4,4_) and at the same time increasing fecundity and the probability of birds transitioning from having a successful fledging event (F) to failing to fledge a chick (stage E, or H) increases population entropy most (Table 4). Reducing the transition between F and H (*p*_3,4_, and increasing fecundity, *p*_1,4_, the transitions to stage E, *p*_2,4_ and stasis of stage F, *p*_4,4_) reduces population entropy most (Table 4).

**Table 4:**
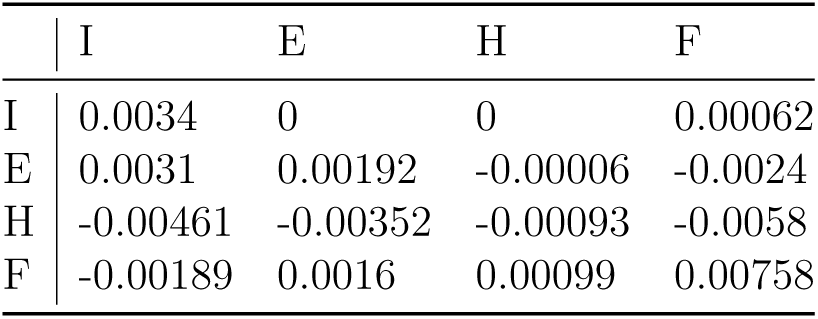
Integrated sensitivity to entropy.

|  | I | E | H | F |
| --- | --- | --- | --- | --- |
| I | 0.0034 | 0 | 0 | 0.00062 |
| E | 0.0031 | 0.00192 | -0.00006 | -0.0024 |
| H | -0.00461 | -0.00352 | -0.00093 | -0.0058 |
| F | -0.00189 | 0.0016 | 0.00099 | 0.00758 |

**Table 5:** Integrated sensitivity to λ.

|  | I | E | H | F |
| --- | --- | --- | --- | --- |
| I | 0.00219 | 0 | 0 | 0.0024 |
| E | -0.00043 | 0.00026 | 0.00007 | -0.00047 |
| H | -0.00022 | 0.00044 | 0.00012 | -0.00024 |
| F | -0.00154 | -0.0007 | -0.00019 | -0.00169 |

These integrated sensitivities of population entropy are distinct from integrated sensitivities with respect to λ. Sensitivities of population entropy quantifies the change in diversification among life course trajectories (Table 4), while sensitivities of population growth quantify the change in fitness (Table 5). Reducing the probability of staying in stage I in consecutive years (I to I transition, *p*_1,1_) and increasing the chance of recruiting to the breeding cohort (I to E,H,F transitions, *p*_2,1_ to *p*_4,1_) increases population growth rate most strongly (Table 5), but is not as influential on diversification of life courses (Table 4). Reducing fecundity (F to I transitions, *p*_1,4_) also leads to a strong increase in population growth when at the same time transitions between F and E, H, F (*p*_2,4_ to *p*_4,4_) are increased (Table 5). The most negative effect for population growth rates are achieved if transitions between I and F (*p*_4,1_), and F and F (*p*_4,4_) are reduced (and at the same time transitions to the other stages are increased, Table 5). This latter observation is not surprising given that we find the highest classical (non-integrated) sensitivities with respect to λ for the same transitions (*p*_4,1_ and *p*_4,4_, Table 3).

The integrated sensitivities with respect to population entropy (Table 4), show which transitions are most critical for generating diversity among life course trajectories, but they do not provide information on whether such diversity might be adaptive or neutral. This understanding, whether diversification of life courses is adaptive or neutral, might not only be informative on a fundamental question in biology, how heterogeneity among individuals evolves and can be maintained, it might also inform on adaptive strategies of niche differentiation expressed as diversification in life courses. The integrated sensitivities with respect to λ (Table 5) provide us with information how changes in transitions affect population growth and fitness. Sensitivities with respect to λ (Table 3) have been used to quantify forces of selection acting on transition probabilities (Caswell, 2001). The higher the sensitivity with respect to λ, the stronger selection should have acted on these transition rates. The integrated sensitivities with respect to λ we compute in Table 5, do not inform us on diversity among life courses. Therefore, to approach the question whether the diversification in life course trajectories measured as the population entropy, might be adaptive, we correlated the two measures of integrated sensitivities for each matrix element. As we see in Fig. 1, the two measures of sensitivity are not correlated and hence the elements that contribute most to diversification of life courses are not those that are under the strongest selection. We also do not find evidence for negative correlation, that is, selections seems not to act against diversification. This suggests that the resulting diversity among life courses might rather be neutral. Such interpretation supports neutral theories of life history evolution (Tuljapurkar et al., 2009; Steiner and Tuljapurkar, 2012), and challenges adaptive theories arguing that variability in life courses is adaptive, an interpretation found in various evolutionary ecological studies (Stearns, 1992). However, our interpretation must be approached with caution since we only explored one of many solutions for the constraints among transition probabilities.

**Figure 1:**
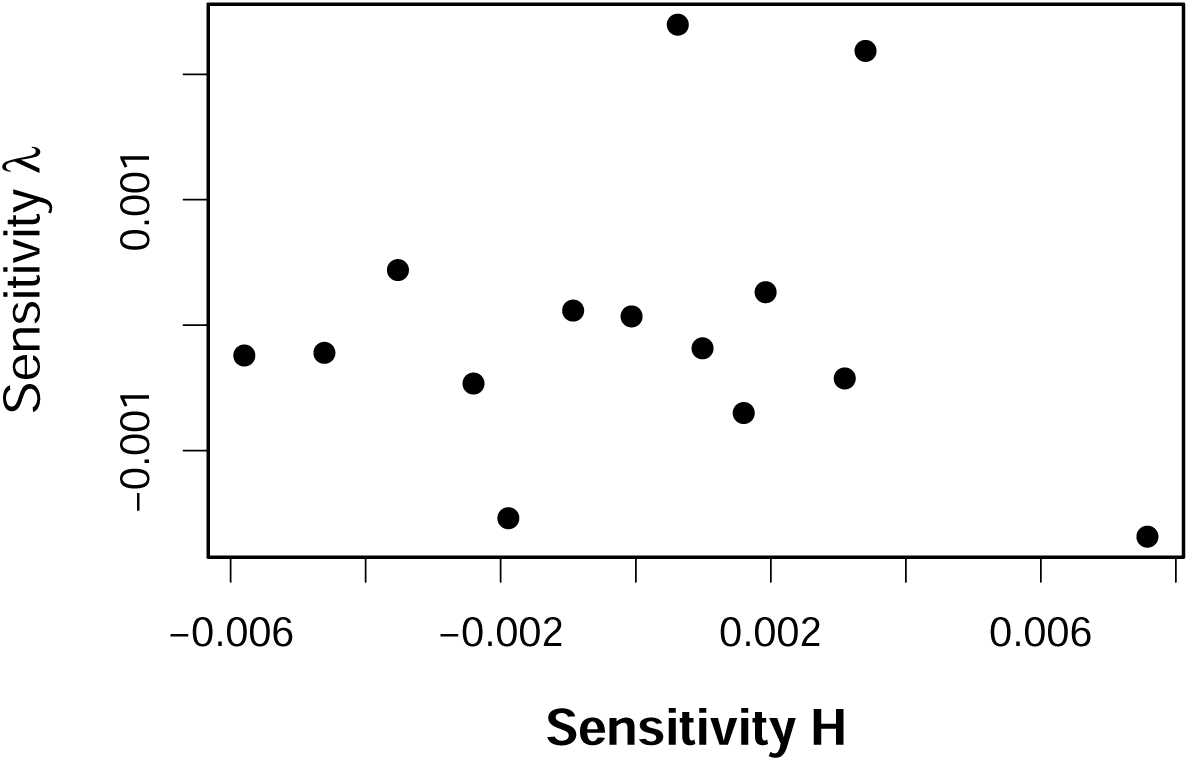
Correlation between sensitivity of entropy and sensitivity of λ.

Our example on the Thick-billed Murre, illustrates how sensitivities of population entropy can be used to approach questions about adaptive diversification in individuals life courses, but our example is only limited to one population. For a more general understanding more species and more solutions to constraints among transition probabilities should be explored.

Population entropy varies substantially among populations and species (Tuljapurkar et al., 2009). Within populations population entropy varies among years, i.e. with varying environments, but the selective forces that shape heterogeneity among individual life courses do not correlate with well-known classical ecological selective forces such as population density (Hernandez-Pacheco and Steiner, 2017). Population entropy also changes with age within a population, indicating changes in transition probabilities with age (Plard et al., 2012). This knowledge on other species and populations show that entropy, as well as fitness varies among populations and conditions experienced by populations. In our example we averaged across environments and across age for simplification and better illustration of the method, but such additional environmental and demographic dimensions can easily be explored. Our motivation to derive the sensitivity with respect to population entropy was mainly to explore the potential evolution of individual stage dynamics, and its effect beyond genotypic, environmental and gene-by-environment interactions. One could ask a different question with a simpler approach: are populations that diversify fast in their life courses more fit? To answer this question one could simply correlate the population entropy to the population growth rate, λ, i.e. one would not use the derivatives (sensitivities to each matrix element) but the population level measure of entropy and growth. These population level demographic parameters do not reveal the influence of the individual stage transitions and which stage transitions contribute most to diversification and fitness. However, the latter information might be crucial to better understand and infer on the underlying mechanisms and allow to go beyond decomposing variance explained by genotypes, environments and their interactions. These insights might also be informative for managing populations and species conservation.

We also like to highlight that neutral and adaptive processes have shaped the transition rates in the stage structured matrix. From a theoretical perspective two matrices with the same population growth rate, can differ vastly in their population entropy, from complete determinism of life courses to maximum entropy (all transition probabilities are equal). Similarly, we can construct matrices that have the same population entropy but differ substantially in their fitness, λ. Such differences are also observed in nature — though perhaps not to the same extreme. For instance, in a free-living monkey population where individuals are closely tracked, heterogeneous trajectories with individuals frequently changing among stages can lead to very similar population structure as can a few trajectories with low level of dynamics, only depending on the environment (Hernandez-Pacheco and Steiner, 2017). The population level stage frequencies do not reveal the underlying differences in individual level stage dynamics. We believe it therefore to be crucial to explore individual level dynamics to understand how diversity in phenotypes and life courses is generated and maintained.

## Conclusions

The sensitivities of the population entropy we derived reveal the transitions among life stages that contribute most to the diversification in life course trajectories (Table 4). We can use these sensitivities of the population entropy in combination with sensitivities on fitness to inform a larger debate on potential selective forces acting on the dynamics and diversification of life courses (Shefferson, 2010). Our example on the Thick-billed Murres illustrates that we only have a limited understanding about changes that generate differences between individuals. In our example the transitions that generate diversity in life courses are not linked to the most sensitive transitions influencing population growth and hence suggest that observed diversification in life courses are neutral rather than adaptive. We have to be cautious about over interpretation of this result, since many solutions for the constraints among transition probabilities exist (Caswell, 2001) and we only have explored one, that seemed to us biologically plausible. Identifying influential stage transitions may not directly reveal the underlying mechanisms that generate diversification but may nonetheless be useful. Mechanistic insights should be easier for populations in which individual stages are closely associated with known underlying mechanisms, for instance via gene expression or methylation. If stages are defined as geographic location, identifying the transitions (migration among locations) that generate most diversification (sensitivity with respect to population entropy) and those that are associated with the highest increase of fitness (sensitivity with respect to λ), might inform niche differentiation and dynamics in metapopulations, and so guide conservation decisions.

## Acknowledgement

We thank Hal Caswell and Troy Day for helpful comments on an early draft.

## A Appendix

### A.1 The Ergodic Case

The perturbation matrix **B** satisfies (1) of the main text. Writing

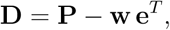

see also that

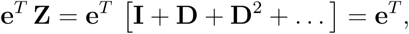

so finally, from (5),

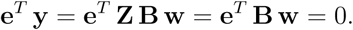

The perturbation of the entropy in (1) uses the expansion

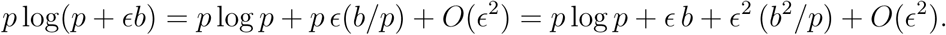

Keeping terms to *O*(*ϵ*) yield three terms (omitting the summations over *I* and *j*),

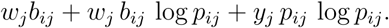

Recall that Σ_*i*_ *b*_*ij*_ = 0 for every *j* to see that the first term is zero, leaving us with equation (8).

### A.2 Conditional Entropy

#### A.2.1 Simplifying the Entropy

The entropy is defined by the middle line of (13). Insert (12) to obtain

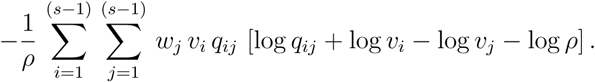

Now use the facts Σ_*i*_ *q*_*ij*_*w*_*j*_ = *ρw*_*i*_,Σ_*i*_ *v*_*i*_*q*_*ij*_ = *ρv*_*j*_ to see that the two middle terms cancel, and to see that the last term (with sums) is just log *ρ*. This yields the last line of equation (13).

#### A.2.2 Perturbing Eigenvectors

We derive (17); proceed similarly to get (18). Now the perturbed right eigenvector of **Q** satisfies the usual equation

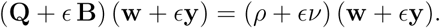

The order *ϵ* terms here are:

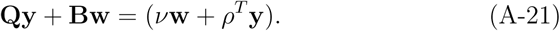

Now note that **w v**^*T*^ is a matrix that projects any vector onto **w**. When we perturb the matrix **Q**, the change **y** must be orthogonal to **w** (otherwise we are just making a proportional change in every matrix element). Hence we must have

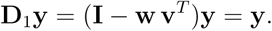

Also

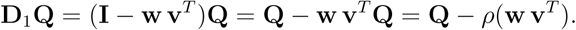

Using these facts, multiply all terms of (A-21) by matrix **D**1 to get, first,

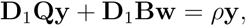

and then

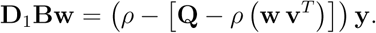

Using the inverse of the matrix on the right (guaranteed to exist because *ρ* is the dominant eigenvalue) leads to (17).

#### A.2.3 Sensitivity of Entropy

We examine separately the two terms of (13) and find perturbations to order *ϵ*. The first term changes to

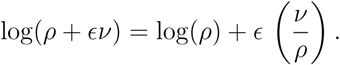

The second term of (13) has the form

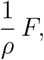

say, where *F* stands for the double sum.

Now (much as in Section A.1) the perturbation of the double sum in (13) is

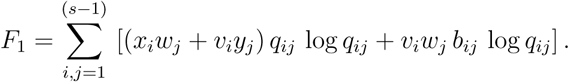

Thus the effect of the perturbation on the second term of (13) is to produce

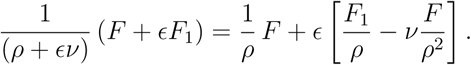

So the total perturbation is

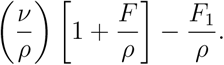

Using (13) to express *F/ρ* in terms of the entropy *H*(**Q**) yields (20).

### A.3 Transforming projection matrix to Markov chain

To transform a population projection model into a Markov chain, we follow Tuljapurkar’s approach (Tuljapurkar, 1982). Note, Tuljapurkar’s projection matrix describes transitions from row to column, whereas our matrix **P** describes transitions from columns to rows, hence the transformation for our matrix is as follows:

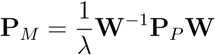

with **P**_*M*_ being the Markov chain (Table 2), **P**_*P*_ being the population projection matrix (Table 1), λ being the population growth rate (dominant eigenvalue of **P**_*P*_), and **W** being a matrix of zeros except for the diagonal elements of (*w*_*i*_), which are the normalized stable stage distribution values (normalized right eigenvector corresponding to dominant eigenvalue of matrix **P**_*P*_). **W**^−1^ is the inverse of matrix **W**.

#### Special Case: Perturbing a Maximum Entropy chain

A chain with maximum entropy has transition matrix elements *p*_*ij*_ = (1*/s*) where, as before, *s* is the number of stages (Tuljapurkar et al., 2009). Clearly **w** has every element equal to (1*/s*) and we can write

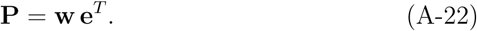

The entropy of this chain is just *H* = log *s* (see also Tuljapurkar et al. (2009)). The chain’s fundamental matrix (see (3)) is just **Z** = **I**, which means that when we perturb the chain to **P** + *ϵ* **B** the eigenvector **w** becomes (see (5)) just **w** + *ϵ* **y** with **y** = **Bw**. The second-order perturbation of **w** is zero (i.e., **y**_2_ = 0).

The sensitivity of this chain is zero! To see that this is true in our equations, observe that in (9) we have

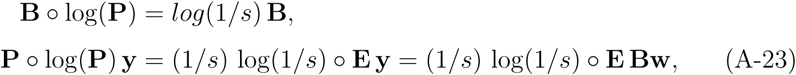

where **E** is a matrix with all elements equal 1. Hence both terms in *H*_1_ (9) are proportional to **e**^*T*^ **B** – but this has to be zero for any possible perturbation (recall the column sums of **B** equal zero), so *H*_1_ = 0. More generally, sensitivity is just a (complicated) derivative of entropy and since we start with maximum entropy it must be true that any derivative of the entropy is zero (that’s what defines a maximum).

So what about *H*_2_ in (10)? Note that here *by*_2_ = 0, and that the arguments in (A-23) imply that the only surviving term in (10) is

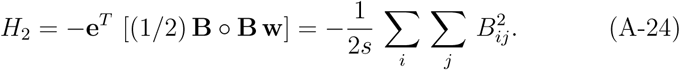

Thus perturbing a maximum entropy chain with transition matrix **P** by the constrained matrix *ϵ* **B** always yields a reduced entropy

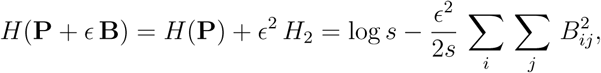

to order *ϵ*^3^.

### A.4 The thick-billed Murre, population projection model

We used data from a total of 1984 individuals, of which 1128 individuals where banded as chicks (immatures), and 856 were banded as adults (left censored). In the breeding colony on Coats Island, these birds were observed over a breeding season and many sightings of uniquely banded bird were made each year. Birds are highly philopatric to their breeding sites which makes it relatively easy to record the breeding outcome for a given year (Steiner and Gaston, 2005). We used 5956 records of annual breeding outcomes of which 1313 were birds laid an egg but not manage to hatch a chick, E; 518 hatch a chick but did not manage to fledge the chick, H, 3031 birds that successfully fledged a chick, F, and 1094 unknown events, U. Since birds are highly philopatric to their breeding site we could assign each bird to a breeding plot. For a few birds that switched a breeding plot within their lifetime, we assigned them to the breeding plot they spent most time breeding at.

The colony on Coats Island is divided into different study plots, and we only included data from six study plots (D, J, K, N, Q, S) that had longitudinal data on a larger number of individuals. For the 1128 immature individuals that were banded as chicks in the colony and then later recruited as breeders, we assumed that they would stay as immatures for the first three years, before they would be allowed to start transitioning to and among the breeding stages (E, H, F, U). Only three of these 1128 birds recruited at age two into the breeding cohort, for these three birds we considered their observed breeding status at age three. Once a bird left the immature stage it was not allowed to transition back to the immature stage. Entering the immature stage from a breeding stage (E, H, F, U) was only possible as a newborn, that is through fertility (Table 1).

Recapture (sighting) effort varied among study plots and years. We therefore accounted for this varying effort among plots and years when we estimated the stage-specific survival and transition probabilities for which we used program MARK (White and Burnham, 1999). This means we accounted for plot and year specific recapture probabilities (mean= 0.41 ± 0.17 Stdev) but not stage-specific recapture probabilities (i.e. we assumed that E,H,F stages are equally likely being sighted). Accounting for these biases ascertained that the probability of a bird surviving or transitioning among stages did not depend on the study plot it bred at, but on its current stage.

Banding of chicks started in 1981 but band reading (sightings) only began in 1991, so all recapture (sighting) probabilities for all plots prior to 1991 were set to 0. Similarly no sighting effort was made for plot D in 2001; for plot J prior to 1995, and in 2000, 2005-2008, 2010, 2011; for plot K in 2001, 2003-2006, and 2011; for plot N in 2001, 2003-2006, and 2011; for plot S in 2000-2002, 2004, 2006, and 2011. In those years for these plots sighting probabilities for the breeding stages (E, H, F, U) were set to 0. For plot Q we had sighting records for each year between 1991 and 2011 and estimated plot specific sighting rates for each year. We did not estimate stage-specific sighting probabilities, but only plot-and year-specific sighting probabilities, since the sighting probability should not depend on the breeding stage (recall we have many observation of each individual within a breeding season).

The data only included birds that recruited as breeders (or attempted breeders) to the colony, we therefore adjusted the immature survival for the population projection model using a previously described estimate of 40.5% of fledglings survival to age three, the age when many individuals started to recruit as breeders (Gaston et al., 1994). This resulted in an annual immature survival of 0.74. Survival rates of the other stages (after correcting for the unknown events) equalled 0.96 for E, 0.87 for H, and 0.91 for F. Table 1 shows the population projection matrix, summarizing the stage transition and survival rates (column sums). The corresponding transformed Markov chain is shown in Table 2.

When we estimated the stage specific transition parameters, using program MARK, we used a multinomial logit function to assure that the transition rates of a given stage sum to 1. This estimation of the survival and transition probabilities included unknown breeding outcomes, U. To account for these unknown breeding outcomes we corrected the survival and transition probabilities of the known breeding stages (E, H, F). We did this by first estimating the fractions of the known breeding outcomes (1313 E, 518 H, 3031 F; i.e. 0.27% E, 0.11% H; and 0.62% F). The expected number of unknown events and their associated survival rates compared to the known events was then taken into account to correct the survival rates of the known stages.

Transition rates to the unknown stage were added to the transition rates of the known stages (E, H, F). We did this by taking the estimated transition probability of a given stage to the unknown stage, and increased each stage transition of the observed stages by its relative weight. This correction was done for each stage (I, E, H, F) and provided the four by four matrix that contributes to Table (1).

Survival estimates of the immature stage, I, based on the MARK model was very close to 1 (if we forced it to be exactly one we had convergence issues). Such a high survival rate is expected since only birds entered the data base if they were recorded as breeders (or attempted breeders), i.e. they all needed to survive the immature stage. In order to get a more realistic population projection model, we reduced annual immature survival to 0.74 which leads to a survival between fledging and age three of 40.5 %; a survival rate reported by Gaston et al. (1994) for this population.

Murres lay a single egg and do not have multiple broods, for that any successful fledgling (stage F event) contributed to fertility. We did only consider female fledglings, assuming that 50% of all fledglings are females. So our resulting population projection model can be seen as a one sex (female) model even though we used male and female observations for estimating survival, transition and sighting probabilities. Other than a slight delay in onset of breeding for males, transition and survival rates have been estimated to be very similar in this species (Gaston et al., 1994). If we only had used data from known females the amount of data would have been much lower and parameter estimations less accurate.

## Notes

#### Summary of Updates

Substantial editing and revisions have been undertaken to clarify the approach.

## References

Barton, N. and Keightley, P. (2002). Understanding quantitative genetic variation. NATURE REVIEWS GENETICS, 3(1):11–21.

Barton, N. H., Etheridge, A. M., and Veber, A. (2017). The infinitesimal model: Definition, derivation, and implications. THEORETICAL POPULATION BIOLOGY, 118:50–73.

Caswell, H. (2001). Matrix population models: construction, analysis, and interpretation, volume 2nd. Sinauer Associates.

Caswell, H. (2009). Stage, age and individual stochasticity in demography. Oikos, 118(12):1763–1782.

Champagnat, N., Ferriere, R., and Meleard, S. (2006). Unifying evolutionary dynamics: From individual stochastic processes to macroscopic models. THEORETICAL POPULATION BIOLOGY, 69(3):297–321.

Darroch, J. N. and Seneta, E. (1967). On Quasi-Stationary Distributions in Absorbing Continuous-Time Finite Markov Chains. Journal of Applied Probability, 4(1):192.

Der, R., Epstein, C. L., and Plotkin, J. B. (2011). Generalized population models and the nature of genetic drift. THEORETICAL POPULATION BIOLOGY, 80(2):80–99.

Diekmann, O., Gyllenberg, M., and Metz, J. (2003). Steady-state analysis of structured population models. THEORETICAL POPULATION BIOLOGY, 63(4):309–338.

Endler, J. A. (1986). Natural selection in the wild, volume 21 of Monographs in Population Biology 21. Princeton University Press.

Evans, S. N. and Steinsaltz, D. (2007). Damage segregation at fissioning may increase growth rates: a superprocess model. Theoretical population biology, 71(4):473–90.

Finch, C. and Kirkwood, T. B. (2000). Chance, Development, and Aging. Oxford University Press, Oxford.

Gaston, A. J., de Forest, L. N., Donaldson, G., and Noble, D. G. (1994). Population Parameters of Thick-Billed Murres at Coats Island, Northwest Territories, Canada. The Condor, 96(4):935–948.

Geoghegan, J. L. and Spencer, H. G. (2012). Population-epigenetic models of selection. THEORETICAL POPULATION BIOLOGY, 81(3):232–242.

Gould, S. and Lewontin, R. (1979). SPANDRELS OF SAN-MARCO AND THE PANGLOSSIAN PARADIGM - A CRITIQUE OF THE ADAPTATIONIST PROGRAM. PROCEEDINGS OF THE ROYAL SOCIETY SERIES B-BIOLOGICAL SCIENCES, 205(1161):581–598.

Hartl, D. J. and Clark, A. (2007). Principles of population genetics. Sinauer, Sunderland.

Hernandez-Pacheco, R. and Steiner, U. K. (2017). Drivers of diversification in individual life courses. The American Naturalist, 190(6):E132–E144.

Hill, M. F., Witman, J. D., and Caswell, H. (2004). Markov chain analysis of succession in a rocky subtidal community. The American naturalist, 164(2):E46–61.

Jouvet, L., Rodriguez-Rojas, A., and Steiner, U. K. (2018). Demographic variability and heterogeneity among individuals within and among clonal bacteria strains. Oikos, 127(5):728–737.

Kato, T. (1966). Perturbation theory for linear operators. Grundlehren der mathematischen Wissenschaften, 132.

Lande, R., Engen, S., and Saether, B. (2003). Stochastic population dynamics in ecology and conservation.

Lenormand, T., Roze, D., and Rousset, F. (2009). Stochasticity in evolution. Trends in ecology & evolution, 24(3):157–165.

Mackay, T. F. C., Stone, E. A., and Ayroles, J. F. (2009). The genetics of quantitative traits: challenges and prospects. NATURE REVIEWS GENETICS, 10(8):565–577.

Matthews, J. P. (1970). A Central Limit Theorem for Absorbing Markov Chains. Biometrika, 57(1):129.

Melbourne, B. a. and Hastings, A. (2008). Extinction risk depends strongly on factors contributing to stochasticity. Nature, 454(7200):100–3.

Ohta, T. and Gillespie, J. (1996). Development of Neutral and Nearly Neutral Theories. Theoretical population biology, 49(2):128–42.

Orr, H. (2005). The genetic theory of adaptation: A brief history. NATURE REVIEWS GENETICS, 6(2):119–127.

Pelletier, F., Garant, D., and Hendry, A. P. (2009). Eco-evolutionary dynamics. PHILOSOPHICAL TRANSACTIONS OF THE ROYAL SOCIETY B-BIOLOGICAL SCIENCES, 364(1523):1483–1489.

Plard, F., Bonenfant, C., Delorme, D., and Gaillard, J. (2012). Modeling reproductive trajectories of roe deer females: Fixed or dynamic heterogeneity? Theoretical Population Biology, 82(4):317–328.

Roze, D. and Rousset, F. (2008). Multilocus models in the infinite island model of population structure. THEORETICAL POPULATION BIOLOGY, 73(4):529–542.

Schweitzer, P. J. (1968). Perturbation Theory and Finite Markov Chains. Journal of Applied Probability, 5(2):401.

Shannon, C. E. (1948). A mathematical theory of communication. Bell system technical journal, 27(3):379–423.

Shefferson, R. (2010). Why are life histories so variable. Nature Education Knowledge, 1(12):1.

Stearns, S. C. (1992). The evolution of life-histories. Oxford University Press, Oxford.

Steiner, U. K. and Gaston, A. (2005). Reproductive consequences of natal dispersal in a highly philopatric seabird. Behavioral Ecology, 16(3):634–639.

Steiner, U. K., Lenart, A., Ni, M., Chen, P., Song, X., Taddei, F., Lindner, A., and Vaupel, J. (2019). Two stochastic processes shape diverse senescence patterns in a single-cell organism. Evolution, page 105387.

Steiner, U. K. and Tuljapurkar, S. (2012). Neutral theory for life histories and individual variability in fitness components. Proceedings of the National Academy of Sciences of the United States of America, 109(12):4684–9.

Steiner, U. K., Tuljapurkar, S., Coulson, T., and Horvitz, C. (2012). Trading stages: life expectancies in structured populations. Experimental gerontology, 47(10):773–81.

Tuljapurkar, S., Steiner, U. K., and Orzack, S. H. (2009). Dynamic heterogeneity in life histories. Ecology letters, 12(1):93–106.

Tuljapurkar, S. D. (1982). Why use population entropy? It determines the rate of convergence. Journal of Mathematical Biology, 13(3):325–337.

Van Tienderen, P. H. (1995). Life cycle trade-offs in matrix population models. Ecology, 76(8):2482–2489.

Vuilleumier, S., Goudet, J., and Perrin, N. (2010). Evolution in heterogeneous populations From migration models to fixation probabilities. THE-ORETICAL POPULATION BIOLOGY, 78(4):250–258.

Walsh, B. (2001). Quantitative genetics in the age of genomics. THEORETICAL POPULATION BIOLOGY, 59(3):175–184.

White, G. C. and Burnham, K. P. (1999). Program MARK: survival estimation from populations of marked animals. Bird Study, 46(sup1):S120–S139.

